# Genome specialization and decay of the strangles pathogen, *Streptococcus equi*, is driven by persistent infection

**DOI:** 10.1101/014118

**Authors:** Simon R. Harris, Carl Robinson, Karen F. Steward, Katy S. Webb, Romain Paillot, Julian Parkhill, Matthew T. G. Holden, Andrew S. Waller

## Abstract

Strangles, the most frequently diagnosed infectious disease of horses worldwide, is caused by *Streptococcus equi*. Despite its prevalence, the global diversity and mechanisms underlying the evolution of *S. equi* as a host-restricted pathogen remain poorly understood. Here we define the global population structure of this important pathogen and reveal a population replacement in the late 19^th^ or early 20^th^ century, contemporaneous with a spate of global conflicts. Our data reveal a dynamic genome that continues to mutate and decay, but also to amplify and acquire genes despite the organism having lost its natural competence and become host-restricted.

The lifestyle of *S. equi* within the horse is defined by short-term acute disease, strangles, followed by long-term carriage. Population analysis reveals evidence of convergent evolution in isolates from post-acute disease samples, as a result of niche adaptation to persistent carriage within a host. Mutations that lead to metabolic streamlining and the loss of virulence determinants are more frequently found in carriage isolates, suggesting that the pathogenic potential of *S. equi* reduces as a consequence of long term residency within the horse post acute disease. An example of this is the deletion of the equibactin siderophore locus that is associated with iron acquisition, which occurs exclusively in carrier isolates, and renders *S. equi* significantly less able to cause acute disease in the natural host. We identify several loci that may similarly be required for the full virulence of *S. equi*, directing future research towards the development of new vaccines against this host-restricted pathogen.

## INTRODUCTION

Bacterial pathogens live a double life, balancing the requirements of acute disease with persistence and colonisation in order to optimise their ability to transmit to naïve animals. These high and low virulence states evolve over time to fit with each host species. However, the impact of evolution towards a persistent state on streamlining the genome of a virulent organism has not been described. Strangles, caused by the host adapted Lancefield group C bacterium *Streptococcus equi* subspecies *equi (S. equi)*, is one of the oldest recognized infectious diseases of horses and continues to cause significant welfare and economic costs throughout the world. The clinical signs of strangles, typified by pyrexia, followed by abscessation of lymph nodes in the head and neck, were first reported by Jordanus Ruffus in 1251 (Rufus 1251). Rupture of lymph node abscesses releases highly infectious pus that can spread the infection from one horse to another, but this in itself cannot explain the global success of *S. equi*. Rather, it is believed that incomplete drainage of abscess material from the retropharyngeal lymph nodes permits the organism to persistently infect the adjacent guttural pouches of ‘carrier’ horses, usually by residing within dried balls of pus known as chondroids (Newton, Wood, Dunn, DeBrauwere and Chanter 1997). *S. equi* can persist in this low nutrient state in the absence of clinical signs for several years and potentially the remaining lifetime of the horse, providing the organism with prolonged opportunity to be shed into the environment and transmit to naïve animals (Newton, Verheyen, Talbot, Timoney, Wood, Lakhani and Chanter 2000). The disease cycle of acute disease followed by persistent infection has underpinned the success of *S. equi*, balancing the requirements of both acute and carrier states.

The worldwide population of *S. equi* is almost clonal by multilocus sequence typing (MLST) (Webb, Jolley, Mitchell, Robinson, Newton, Maiden and Waller 2008), consisting of only two STs, ST179 and ST151, and has been proposed to have evolved recently from an ancestral strain of *Streptococcus equi* subspecies *zooepidemicus (S. zooepidemicus)* (Webb, Jolley, Mitchell, Robinson, Newton, Maiden and Waller 2008). Both *S. equi* and *S. zooepidemicus* belong to the same group of pyogenic streptococci as the important human pathogen *Streptococcus pyogenes* (Group A *Streptococcus)* and share many common virulence mechanisms, with evidence of cross-species horizontal DNA exchange (Holden, Heather, Paillot, Steward, Webb, Ainslie, Jourdan, Bason, Holroyd, Mungall et al. 2009; Lefebure, Richards, Lang, Pavinski-Bitar and Stanhope 2012). Comparison of the complete genomes of *S. equi* strain 4047 (Se4047) and *S. zooepidemicus* strain H70 (SzH70) showed that the pathogenic specialization of *S. equi* has been shaped by a combination of gene loss due to nonsense mutations and deletions, and gene gain through the acquisition of mobile genetic elements (MGEs) (Holden, Heather, Paillot, Steward, Webb, Ainslie, Jourdan, Bason, Holroyd, Mungall et al. 2009). However, the selective forces underlying this evolutionary process as the pathogen transitions between acute disease and persistent carriage remain unknown.

In this study we utilize whole genome sequencing of a large collection of *S. equi* strains from around the globe. The collection contains isolates from horses that were part of outbreaks on large farms in the UK, including isolates from acute and persistent carriage states, and also multiple isolates from the same animal. Using these data we investigate the evolutionary history of the species, and uncover evidence of common pathways of within-host adaptation associated with persistent carriage and social evolution.

## RESULTS

### Population structure of a global collection of *S. equi*

The genomes of a collection of 224 *S. equi* isolates were sequenced (Supplementary Table 1). Forty geographically and temporally diverse isolates from outbreaks in Australia, Belgium, Canada, Ireland, New Zealand, Saudi Arabia, Sweden, and the USA were included to provide a global snapshot of the diversity of *S. equi*. Allied with this, 180 acute infection and carriage isolates from 41 counties across the UK were included to determine the diversity of *S. equi* within individual outbreaks. Finally, isolates from commercially available live attenuated vaccines, Equilis StrepE (Jacobs, Goovaerts, Nuijten, Theelen, Hartford and Foster 2000) and Pinnacle IN (high and low capsule) (Walker and Timoney 2002), and the Se1866 isolate, which is the basis of the new multi-component Strangvac vaccine (Guss, Flock, Frykberg, Waller, Robinson, Smith and Flock 2009), were included to determine the relationships of these vaccines to the currently circulating population of *S. equi*, and to identify mutations responsible for attenuation of the Pinnacle IN strain.

The indexed sequencing reads from each isolate were mapped against the *S. equi* Se4047 reference genome (Holden, Heather, Paillot, Steward, Webb, Ainslie, Jourdan, Bason, Holroyd, Mungall et al. 2009) to identify single-nucleotide polymorphisms (SNPs), insertions and deletions (indels). 3,109 polymorphic sites were identified across the genome. 58.8% of these sites (*n* = 1,844) were in accessory regions of the Se4047 reference genome, which corresponds to only 16.4% of the length of the genome. Much of this variation in accessory regions is likely to have arisen by recombination and horizontal gene transfer of alternative MGEs. To remove confounding signals in these regions, all phylogenetic analyses were carried out using the 1,265 polymorphic sites (Supplementary Table 2) in the 1.9 Mb of the core genome.

A Maximum likelihood (ML) phylogenetic reconstruction of the core variable sites of the sequenced isolates is shown in Fig. 1a. Isolates are colored according to clusters defined using a Bayesian method for subdividing populations based on sequence similarity (Corander, Marttinen, Sirén and Tang 2008). Clusters one to three correspond to the three major lineages in the ML tree, with the remainder of isolates falling into a fourth, polyphyletic cluster. The increasingly prevalent ST151 formed a clade within cluster 1, distant from the Equilis StrepE and Pinnacle IN vaccine samples (Fig. 1a). Two features of the tree were particularly striking. First, the observed sequence diversity was surprisingly low for a global collection of isolates of a pathogen with a historical record extending back to at least the 13^th^ century. Second, the tree includes a number of anomalously long branches towards its tips, which appeared inconsistent with the isolation dates of the samples.

**Figure 1.**
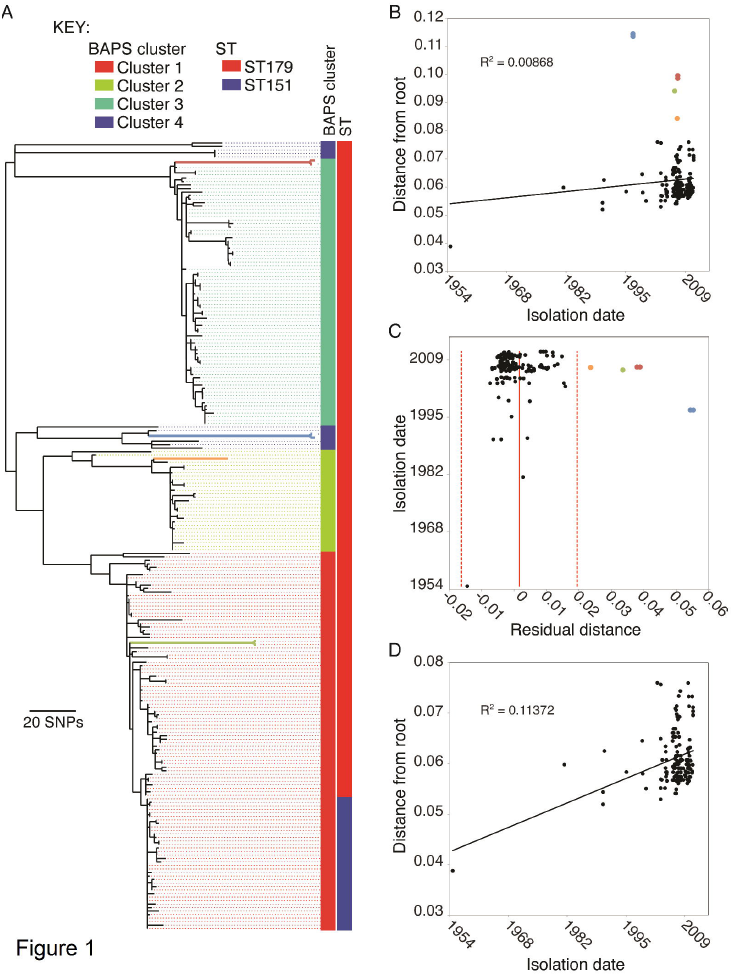
Phylogentic reconstruction and assessment of temporal signal A) Maximum-likelihood phylogeny of *S. equi* based on core genome SNPs. Clusters are colored as reconstructed by BAPs. Columns adjacent to the tree represent the BAPS cluster and MLST type associated with the strains. Branch colors relate branches on the tree to points in panels B and C. B) Root-to-tip plot of all isolates in the ML tree in panel A. C) Residual plot of root-to-tip plot in B. Unbroken red line indicates the mean, and dashed red lines two standard deviations from the mean. D) Root-to-tip plot with the isolates falling outside of the 95% confidence intervals in B excluded.

### Variation in the rate of substitution in the *S. equi* core genome

Plotting root-to-tip distance against isolation date for each isolate for which dating information was available revealed a poor correlation between accumulation of substitutions and time (Fig. 1b) (R^2^ = 0.00868), although a tip-date permutation test (Firth, Kitchen, Shapiro, Suchard, Holmes and Rambaut 2010) confirmed significant temporal signature in the data (*p* = 0.001 from 1,000 permutations). Similar observations in other streptococci have been the result of the effect of homologous recombination, which can import large amounts of variation into the genome *en masse*, leading to elongation of branches and the loss of clock-like mutational signals. Although it has been thought that *S. equi* has lost natural competence (Holden, Heather, Paillot, Steward, Webb, Ainslie, Jourdan, Bason, Holroyd, Mungall et al. 2009), we investigated the possibility that recombination was the cause of the observed long branches. By reconstructing sequences for ancestral nodes in the tree, evidence for past recombination events can be identified in two ways: recombination events from donors outside of the diversity of the tree will cause imports of large numbers of clustered substitutions on individual branches of the tree, while recombination events between donors and recipients within the diversity of the tree will lead to accumulation of homoplasies. In contrast to other streptococci (Cornejo, Lefebure, Bitar, Lang, Richards, Eilertson, Do, Beighton, Zeng, Ahn et al. 2013; Croucher, Harris, Fraser, Quail, Burton, van der Linden, McGee, von Gottberg, Song, Ko et al. 2011; Marttinen, Hanage, Croucher, Connor, Harris, Bentley and Corander 2011), we observed low levels of both SNP clustering (Supplementary Fig. 1) and homoplasy (only 47 or 3.6% of core variable sites), ruling out recombination as the cause of the long branches. The highest density of SNPs (3.2% of SNPs, *p* < 0.0001) and homoplasies (28% of homoplasic SNPs, *p* < 0.0001) in the core genome coincided with the 5’ 1.2 kb region (0.06% of the core genome length) of the of the SeM gene that encodes a fibronectin-binding protein, a virulence factor that binds to fibrinogen and immunoglobulin. The promoter region of the same gene also contained three homoplasies. Similarly, the fibronectin-or fibrinogen-binding proteins, *fneE* and *SzPSe* also exhibit high SNP density and homoplasy, highlighting the importance to *S. equi* of generating variation in these proteins.

Given the observed lack of homologous recombination, the poor root-to-tip correlation must have resulted from variation in substitution rate across the tree. Plotting the residuals for the root-to-tip analysis (Fig. 1c) showed that seven isolates, subtended by four long branches when mapped onto the tree (Fig. 1a), fell outside of two standard deviations of the mean. These included two isolates of the Pinnacle IN vaccine strain; two isolates extracted from the right guttural pouch of a horse (JKS121) sampled during a strangles outbreak in Leicestershire, two isolates from the guttural pouches of a horse (JKS628) sampled during an outbreak in Essex, and one isolate from the right guttural pouch of a horse (JKS731) sampled during an outbreak in Lincolnshire. Excluding these seven isolates increased the root-to-tip R^2^ value to 0.11372 (Fig. 1d). The long branch associated with the Pinnacle IN isolates resulted from 68 shared unique SNPs in the core genome of these isolates. This diversity could be explained by the methods used to make the vaccine, which was a live attenuated vaccine generated by chemical mutagenesis of *S. equi* with N-methyl-N'-nitro-N-nitrosoguanidine (NTG) (Timoney 1985). As expected for NTG-treatment, the substitution spectra for the SNPs unique to the Pinnacle isolates was significantly enriched for C->T and G->A, but deficient in A->G and T->C transitions (Harper and Lee 2012) when compared to other branches in the tree (Supplementary Fig. 2). In wild-type isolates, disruption of the mismatch repair system can also lead to increased substitution rates, but is characterized by an increase in the accumulation of A->G and T->C transitions. We found no significant differences between the substitution spectra of the three long branches leading to clinical isolates, including two isolates from JKS121 which exhibited a nonsynonymous mutation in the mismatch repair gene, *mutS* (Supplementary Fig. 2). Consistent with this, no significant differences (*p* = 0.093) in resistance mutation frequency were found when comparing the long-branch isolates from JKS121, JKS628, JKS731, the Pinnacle IN vaccine isolates and the reference Se4047 *in vitro* (Supplementary Fig. 3).

### Sequencing resolved the complex epidemiology of strangles outbreaks

JKS121, JKS628 and JKS731 were all sampled during strangles outbreaks in the UK. The inclusion of multiple isolates from outbreaks allows us to utilize the resolution that whole genome sequencing provides to conduct detailed epidemiological analysis of strangles. In most cases, outbreak isolates were highly clonal, and differed by only a small number of SNPs (Fig. 1a), consistent with an import from a single source. However, in some cases, including the outbreaks involving JKS121, JKS628 and JKS731, both active strangles strains, and persistent carriage strains from chondroids were identified from horses at the same stables during a strangles outbreak. In one case, 10 isolates recovered from a small outbreak in Essex over a 5 month period all fell into cluster 3 on our phylogeny. However, within this cluster the isolates grouped into three distinct sub-clades differentiating an outbreak strain and two persistent strains (Fig. 2). Interestingly, of the four isolates recovered from JKS628, including the two outliers in our root-to-tip analysis, two fell into each of the persistent clades possibly reflecting separate, long-standing persistent infections prior to the purchase of this horse some 15 months before the outbreak. A similar situation was observed for six isolates from an outbreak in Leicestershire in 2007. Four isolates, from both carriage and disease, formed a clade in cluster 3, representing the outbreak strain. However, the two root-to-tip outliers from JKS121 were distinct, falling in cluster 1. These two carriage isolates possibly originated from a previous outbreak during the 8-year residency of this horse on the affected premises. Finally, JKS731 was sampled during a large outbreak in Lincolnshire that persisted from 2006 to 2008 and affected over 200 horses. Twenty-seven isolates from this outbreak formed a single sub-clade within cluster 2 on our tree. Two of these isolates, including the carriage isolate from JKS731, which was the final outlier in our root-to-tip analysis, shared a much deeper common ancestor than the rest of the outbreak isolates, which suggests these may be persistent isolates from a historic outbreak of a similar genotype, possibly occurring during the 15 years residency of this horse at the affected premises. The outbreak strain may have been the result of reinfection from an external source, or more likely from a persistent carrier at the stables. As with the Essex and Leicestershire outbreaks, isolates taken during the Lincolnshire outbreak also included two persistent lineages that clustered far away from the main outbreak clade, illustrating the prevalence of long-term carriage of *S. equi*.

**Figure 2.**
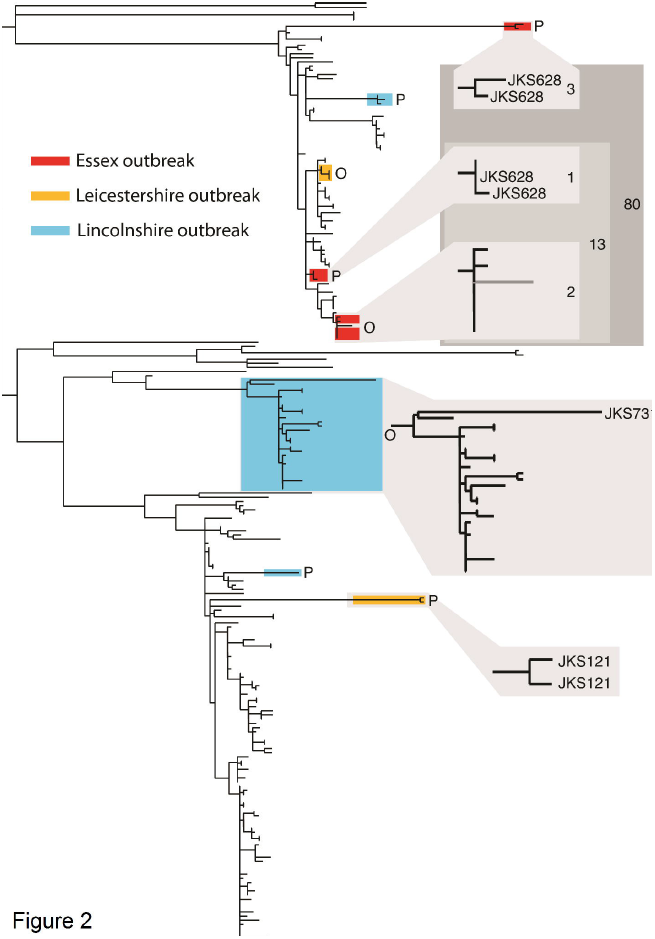
Detailed epidemiology of three *S. equi* outbreaks in the UK. Isolates highlighted on the phylogeny were collected from three stables experiencing strangles outbreaks, as indicated by the key. In each case, isolates did not fall into single clades on the phylogeny, instead being grouped into outbreak (O) and persistent (P) clusters. The positions of the outlier branches identified by root-to-tip analysis are labeled in blow-ups of the relevant clades. For the Essex outbreak, the number of SNPs within and between clades are indicated by the numbers in the grey boxes.

### Substitution rates vary between acute infection and carriage

Remarkably, despite the five clinical isolates identified as outliers by our root-to-tip analysis being sampled during outbreaks, all were isolated from chondroids extracted from guttural pouches and all were identified independently as being persistent isolates representative of historic infection. This observation raised the possibility that the poor temporal signal identified in our root-to-tip analysis was the result of *S. equi* displaying altered substitution rates during persistent infection. To shed further light on this, we analyzed the subset of our dataset for which accurate isolation dates were available with the Bayesian phylogenetics software, BEAST (Drummond and Rambaut 2007a), which allows modeling of molecular clock rates to provide estimations of divergence dates on nodes of a phylogenetic tree. Removal of artificially attenuated vaccine isolates and isolates without dating information produced a dataset of 209 isolates with 1184 polymorphic core genome positions. BEAST includes a number of relaxed molecular clock models which permit modeling of variation in substitution rates on different branches of the tree, allowing us to correct for the observed rate variation in our data, and also to identify other branches exhibiting particularly high substitution rates. Indeed, the combination of a skyline population model and relaxed-exponential clock model was found to be the best fit to our data based on Bayes Factors using the harmonic mean estimator. The topology of the maximum clade credibility (MCC) tree generated from combined data post burn-in from three independent runs of this model was concordant with the ML tree of all isolates (Supplementary Fig. 4). The mean substitution rate per core genome site per year was calculated as 5.22 × 10^-7^ (95% highest posterior density (HPD): 4.04 × 10^-7^ to 6.51 × 10^-7^), slower than the core genome rates reported for many other Gram-positive bacteria, including *Staphylococcus aureus* (3.3 × 10^-6^) (Harris, Feil, Holden, Quail, Nickerson, Chantratita, Gardete, Tavares, Day, Lindsay et al. 2010; Holden, Hsu, Kurt, Weinert, Mather, Harris, Strommenger, Layer, Witte, de Lencastre et al. 2013) and *Streptococcus pneumonia* (1.57 × 10^-6^) (Croucher, Harris, Fraser, Quail, Burton, van der Linden, McGee, von Gottberg, Song, Ko et al. 2011). The analysis provided a median estimate for the time of the most recent common ancestor (tMRCA) of our global sample of *S. equi* to 1909 (95% HPD: 1819 to 1946). Given the historical record of strangles dates back to at least the 13^th^ century, this suggests a global population replacement occurred during the 19^th^ or early 20^th^ centuries, corresponding to a time when horses were a major mode of transport and played important roles in a number of global conflicts (Fig. 3). To ensure this result was not an artifact of the clock or population models employed in our analysis, we calculated the mean, median and 95% HPD tMRCA for a range of model combinations and found that our chosen model exhibited the widest HPD, which encompassed the 95% HPDs of all other model combinations (Fig. 3).

**Figure 3.**
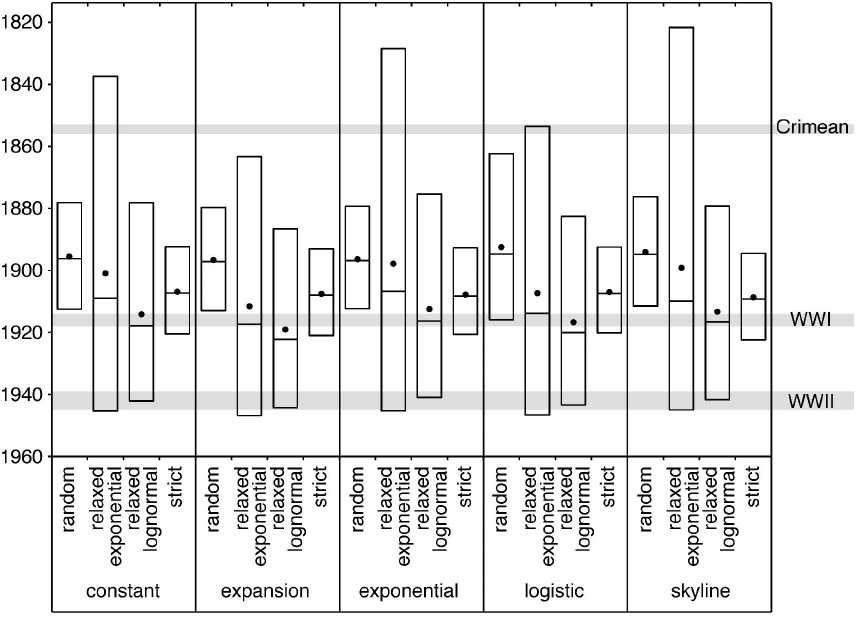
Box plots showing mean (dots), median (center horizontal line) and 95% HPD (highest posterior density) for the time to the most recent common ancestor (tMRCA) of all isolates in the study under various combinations of clock and population model in BEAST. The dates of the Crimean War, WWI and WWI are indicated by the gray shaded regions.

To test our hypothesis that persistent isolates exhibited increased substitution rates, we performed a two-sided Mann-Whitney U test of median substitution rates for branches in our BEAST tree leading to carriage isolates against those leading to isolates from acute infection (Supplementary Fig. 5a). We found a significant difference (p = 0.0004) in rates between the two populations, with persistent isolates exhibiting a significantly increased rate. A linear regression of substitution rate against time for persistent, acute and other (where the site was unknown) branches showed an increasing rate over time for persistent isolates (R^2^ = 20156, *p* = 2.43e^-7^), while the rates for acute (R^2^ = 0.01459, *p* = 0.3693) and unknown (R^2^ = 0.00323, *p* = 0.3815) branches showed no increase with time Supplementary Fig. 5b.

### Variation in the accessory genome was not associated with carriage or infection

Having observed distinct differences in the mutation rates of core genomes of the carriage and acute populations, we undertook a comparative genomic analysis to investigate if there were also differences in the dynamics of the accessory genome. Amongst the mobile elements described in the reference Se4047 isolate, the prophages φSeq2-4, and the integrative conjugative elements ICE*Se1* and ICE*Se2* were the most stable, being almost ubiquitously conserved in the population (Supplementary Fig. 6). In contrast, the prophage φSeq1 appeared more dynamic, with evidence of alternative, but related, prophages in the accessory genomes of isolates belonging to all four clusters. Although the pattern of presence of these prophage elements generally followed the phylogeny, there was evidence of lateral transfer or convergent gain of some prophages in multiple clusters. The deletion of an ancestral CRISPR locus, prior to the emergence of *S. equi* from *S. zooepidemicus*, potentially contributes to the poly-lysogeny of this species (Holden, Heather, Paillot, Steward, Webb, Ainslie, Jourdan, Bason, Holroyd, Mungall et al. 2009).

Although φSeq2-4, ICE *Se1* and ICE*Se2* were generally conserved (Supplementary Fig. 6), we found evidence of both deletions and duplications affecting regions of these elements. Several isolates showed duplications of the phospholipase A_2_ gene, *slaA*, on φSeq2, suggesting that these strains had acquired additional distinct prophages that also contain this putative virulence cargo (Holden, Heather, Paillot, Steward, Webb, Ainslie, Jourdan, Bason, Holroyd, Mungall et al. 2009). Convergent loss of the *seeL* and *seeM* superantigen-encoding genes in six isolates from three distinct lineages is likely to have occurred by horizontal replacement of φSeq3 with alternative phages.

### IS-elements mediate genome decay and gene amplification during carriage

Whilst our comparative analysis revealed that, as would be expected, prophages were mediating the loss and gain of genes, it also revealed that genes were being lost and duplicated in the core genome. The mechanism that appears to be driving the observed deletion and copy-number variation is homologous recombination between insertion sequence (IS) elements. The *S. equi* genome has large expansions of a limited number of IS families, which is hypothesized to result from the evolutionary bottleneck associated with its host-adaptation and pathogenic specialization (Holden, Heather, Paillot, Steward, Webb, Ainslie, Jourdan, Bason, Holroyd, Mungall et al. 2009). We identified an additional 37 novel IS element insertions belonging to existing families across the population, eight of which inserted at the same genomic location in multiple lineages (Supplementary Fig. 7 and 8). IS elements flanked four of the 15 deleted loci and 10 of the 16 amplified regions in the core genome, showing that they play an important role in variation of gene content.

Consistent with our hypothesis, large deletion and duplication events were far more common in carriage isolates than those from acute infection (Supplementary Fig. 7). Within the core genome, the *has* operon, encoding hyaluronic acid capsule biosynthesis components, was the locus most frequently affected by deletions and duplications, with seven independent deletion variants and eight independent amplifications of the locus, which is bordered by IS3 elements (Fig. 4). The same region also exhibited high SNP densities and four independent nonsense mutations. Importantly, the SNPs, deletions and amplifications at this locus were significantly associated with isolates recovered from persistently infected carriers rather than those with acute disease (*p* < 0.0001). Similarly, the citrate cluster, required for the utilization of citrate as a carbon and energy source, was amplified on three separate occasions and deleted twice in carriage isolates. In isolates from acute infection only a single amplification was seen (Supplementary Fig. 7). Somewhat paradoxically, our data suggests that these core metabolic functions are experiencing differential selective pressures within the host, driving both the loss and amplification of these functions during long-term carriage.

**Figure 4.**
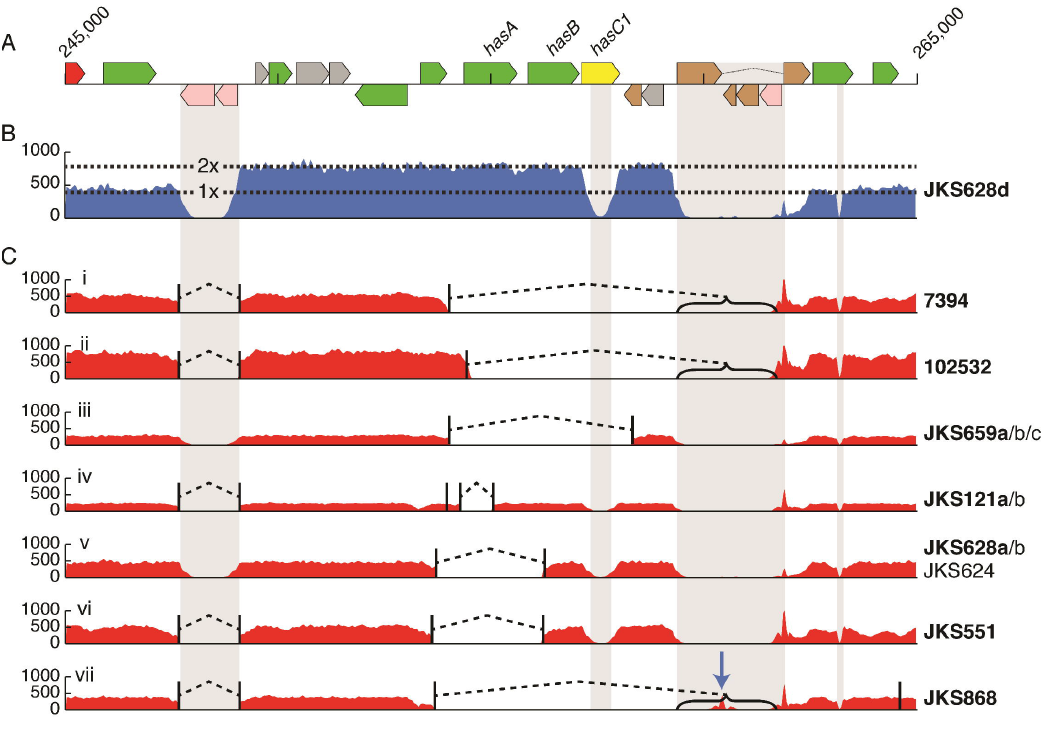
Convergent deletion and amplification of the *has* locus in isolates recovered from independent persistently infected carriers. A) A representation of the genome annotation around the *has* locus. B) Eight isolates, all from different horses, independently exhibited duplications of the region between IS repeats (indicated by gray columns). This plot represents mapped sequence read coverage of JKS628d, showing an increase in coverage spanning the *has* operon, representing duplication of the region. Read mapping reduction in the gray repeat regions is due to an inability to map reads uniquely to those regions. C) 12 isolates from 8 horses exhibited 7 independent deletions of different regions of the *has* locus. Plots i to vii show read mapping of the *has* locus in example isolates for each of the deletions. Names to the right of the plots indicate isolates exhibiting the same deletion. Isolate names in bold are the examples used for the mapping plot. Read mapping drops to zero in the gray repeat regions due to an inability to map reads uniquely to this region. Deletion breakpoints, identified by finding breakpoints in reads, are indicated by vertical black lines joined by dashed bridges. Brackets indicate where one breakpoint location is uncertain due to falling in a repetitive region. The blue arrow in vii indicates the location of a novel IS insertion.

Comparison of the multiple isolates recovered from the same persistently infected horse, JKS628, showed that even these extremely closely related isolates varied by deletions and/or amplifications, demonstrating that microevolution of *S. equi* in the guttural pouch yields a mixture of variant strains, suggestive of social evolution, i.e. the selective pressures on a given cell may be determined by the physiological status of surrounding cells, and therefore heterogeneity in the population drives differential evolution. Isolates JKS628c and JKS628d, recovered from the right and left guttural pouches, respectively, contained a 39.5 kb deletion (SEQ_1232 to SEQ_1253) in ICE*Se2* that included the entire equibactin locus. This locus encodes a putative siderophore, which is known to increase the ability of *S. equi* to acquire iron (Heather, Holden, Steward, Parkhill, Song, Challis, Robinson, Davis-Poynter and Waller 2008), and may enhance survival within, and abscessation of, lymph nodes (Holden, Heather, Paillot, Steward, Webb, Ainslie, Jourdan, Bason, Holroyd, Mungall et al. 2009). This siderophore has been proposed to represent the key speciation event in the evolution of *S. equi* (Holden, Heather, Paillot, Steward, Webb, Ainslie, Jourdan, Bason, Holroyd, Mungall et al. 2009). Three other deletion events (Fig. 5) that are predicted to disrupt equibactin production were identified in isolates recovered from three more, unrelated persistently infected carriers. In contrast, the equibactin region was not deleted in any of the acute isolates (*p* = 0.0528), suggesting that loss of equibactin may preclude transmission to acute infection. To determine if the loss of the equibactin locus represented a dead end mutation that would prohibit these isolates from being able to cause acute disease and therefore continue to be transmitted via the strangles disease cycle, we challenged two groups of seven Welsh mountain ponies with 1 × 10^8^ cfu of Se4047 or an *eqbE* deletion mutant, which is unable to produce equibactin (Heather, Holden, Steward, Parkhill, Song, Challis, Robinson, Davis-Poynter and Waller 2008). Ponies were monitored carefully for early signs of disease, such as pyrexia and a preference for hay over pelleted food, and euthanized before the onset of later clinical signs of strangles (Guss, Flock, Frykberg, Waller, Robinson, Smith and Flock 2009). Deletion of *eqbE* significantly reduced the amount of pyrexia (*p* = 0.021) and pathology at post mortem examination (*p* = 0.041) (Fig. 6). However, loss of *eqbE* did not completely prevent abscess formation with two Δ*eqbE*-infected ponies developing bilateral retropharyngeal lymph node abscesses compared with all seven wild-type infected animals (*p* = 0.021). One Δ*eqbE*-challenged pony developed a submandibular lymph node abscess, two ponies had micro-abscesses without clinical signs of disease and two had no signs of infection. The attenuation of the Δ*eqbE* deletion mutant in Welsh mountain ponies demonstrates that the acquisition of ICE*Se2* was indeed a key step in the evolution of *S. equi* and that strains containing deletions in the equibactin locus, which are shed from persistently infected horses, have a lower virulence potential.

**Figure 5.**
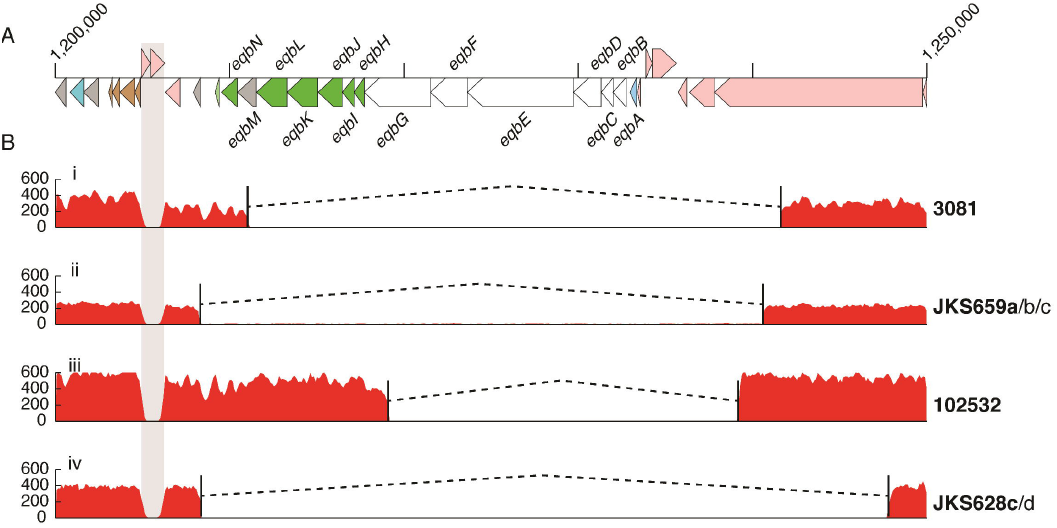
Convergent deletion of the equibactin locus in 7 isolates recovered from four independent persistently infected carriers. A) A representation of the genome annotation around the equibactin cluster. B) Read mapping of the equibactin locus in example isolates exhibiting the four deletion variants. Names to the right of the plots indicate isolates exhibiting the same deletion. Isolate names in bold are the examples used for the mapping plot. Read mapping drops to zero in the gray repeat region due to an inability to map reads uniquely to this region.

**Figure 6.**
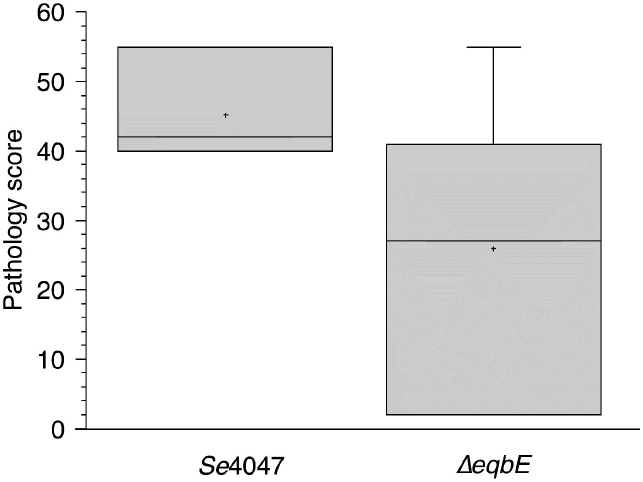
Pathology score of two groups of seven Welsh mountain ponies challenged with wildtype Se4047 or the *ΔeqbE* deletion mutant. The box illustrates the 50% distribution, the line indicates the median value, the cross indicates the mean and error bars indicate the maximum and minimum distribution.

## DISCUSSION

Analysis of a global collection of *S. equi* isolates, has, for the first time, allowed important questions regarding the complex epidemiology of strangles outbreaks to be resolved, and provided compelling evidence of co-infection of individual horses. Variation between isolates in the collection was strikingly low given that strangles was first documented over 700 years ago. Bayesian analysis estimated with 95% confidence that the currently circulating worldwide strains of *S. equi* diverged from a common ancestor between 1819 and 1946. This period of history endured a series of global conflicts including the Crimean War (1853-1856), World War I (WWI, 1914-1918) and WWII (1939-1945) during which horses performed critical roles for all sides (Clabby 1976). Strangles was, and remains, a particular problem in army horses (Bazeley 1940; Todd 1910). Indeed, the Leicestershire outbreak that included horse JKS121 occurred at the Defence Animal Centre in Melton Mowbray. In WWI the British army alone used over a million supply horses, gun horses, cavalry and mules, with at times over 1,000 animals per week being shipped in from the USA to bolster numbers. Despite officers being warned to be on the lookout for acute cases of strangles, the mobilization and mixing of large populations of horses from around the world in an environment ideally suited to the transmission of *S. equi* would have provided the perfect conditions for the mixing of ancestral strains, lateral transfer of mobile genetic elements and the emergence, dissemination and dominance of the fittest strain. At the same time, the dramatic increase in host mortality would have caused repeated transmission bottlenecks for the pathogen, with the associated potential for loss of population diversity. However, the death of eight million horses during WWI would have also removed a significant proportion of host herd immunity.

Government initiatives to increase the supply of high quality light weight cavalry horses, such as the establishment of the National Stud in 1916, produced naïve young horses on an unprecedented scale, providing new hosts for successful *S. equi* clones to colonize. Thus the unique circumstances created by 19^th^ and early 20^th^ century global conflicts would have provided ideal conditions for the emergence of the contemporary population of *S. equi*.

The crucial factor allowing *S. equi* to propagate the disease cycle of strangles is its ability to persist for long periods in a subclinical, but infectious carriage state. The importance of the establishment of persistent infections to the capability of *S. equi* to transmit to naïve animals was first recognized by Captain Todd of the British Army Veterinary Corps as early as 1910 (Todd 1910) and is undoubtedly a major factor in its continued global success. However, it was not until 1997 that the anatomical site of such persistent infections was shown to be the guttural pouch (Newton, Wood, Dunn, DeBrauwere and Chanter 1997). Our data demonstrate that the environment within the guttural pouch, and the chondroid in which *S. equi* persists, drives both the diversification and decay of its genome, and potentially explains the path to host restriction. Although adaptation to the carriage environment, such as loss of the equibactin siderophore, may reduce virulence and transmissibility, leading to an evolutionary dead-end, it is clear that strangles outbreaks can be founded by long-term carriage strains. Indeed, persistence may also bring beneficial effects beyond simply providing time for contact with new naïve hosts. For example, mutation of the antigenic, sortase-processed surface proteins SeM, SzPSe and SEQ_0402 during carriage could assist the evasion of acquired immunity facilitating persistence in the guttural pouch and potentially the recrudescence of strangles. Increased shedding and/or more efficient transmission of new variant strains in this manner provides one explanation for the raised SNP densities observed in the genes encoding these proteins and highlights further genes which exhibit similar raised SNP densities, such as *fneE*, which will direct the design of improved vaccines against strangles. An interesting observation is that the increasingly prevalent ST151 clade of *S. equi* is distantly placed from both the Equilis StrepE and Pinnacle IN vaccine isolates, raising the prospect that this lineage may be escaping current vaccines and should be closely monitored.

The lifecycles of many pathogenic bacteria involve both acute and persistent phases of infection. Balancing the conflicting genetic requirements to cause clinical disease and then to survive in the face of a convalescent immune response represents a significant biological challenge that may ultimately lead towards host-restriction. By sequencing multiple isolates from the same infected horses our analyses have, for the first time, shed light on the complex population dynamics of *S. equi* during carriage within a host. The physiological conditions experienced by individual *S. equi* cells within chondroids will vary according to their immediate environment and the genetic make-up of the local bacterial population with which they interact, generating local selective pressures and driving social evolution. Our data show gene content variation between isolates from long-term carriage *S. equi* populations. Loss of gene function in some cells may be complemented by near-neighbors, or compensated by the physiological and physical benefit of existing in the biological community within chondroids. For example, the production of public goods such as the exopolysaccaride capsule or equibactin siderophore by some cells may compensate for their loss by near-neighbors. The combination of the ability to persist in healthy animals for long periods while remaining infectious and the flexibility to exchange and acquire mobile genetic elements and streamline its genome has played, and will continue to play, a key role in ensuring that *S. equi* remains a persistent threat to equine health with transmission benefitting from the continued mobility of modern horse populations.

## METHODS

### Study collection

Se4047 was isolated from a horse with strangles in the New Forest, Hampshire, UK in 1990. The origins and details of the 224 isolates of *S. equi* that were sequenced in this study are listed in Supplementary Table 1. β-hemolytic colonies of *S. equi* strains were recovered from glycerol stocks following overnight growth on COBA strep select plates (bioMérieux). Their identity was confirmed by a lack of fermentation of trehalose, ribose and sorbitol in Purple broth (Becton Dickinson).

### DNA preparation

A single colony of each *S. equi* strain was grown overnight in 3 ml of Todd Hewitt broth containing 30 μg/ml hyaluronidase (Sigma), centrifuged, the pellet re-suspended in 200 μl Gram +ve lysis solution (GenElute, Sigma) containing 250 units/ml mutanolysin and 2×10^6^ units/ml lysozyme and incubated for 1 hour at 37 °C to allow efficient cell lysis. DNA was then purified using GenElute spin columns according to manufacturer’s instructions (all Sigma).

### DNA Sequencing and variation detection

Library construction for Illumina sequencing was carried out as described previously (Quail, Otto, Gu, Harris, Skelly, McQuillan, Swerdlow and Oyola 2012). One isolate, NCTC9682, was sequenced on an Illumina GAII for 54 cycles from each end, producing paired-end reads with an expected insert size of 250 bp. The remaining isolates were mixed in pools of between 20 and 24 to produce multiplexed libraries that were sequenced on the Illumina HiSeq platform for 75 cycles from each end plus an 8-base index-sequence read. Again the expected insert size between the paired-end reads was 250 bp. Short reads were mapped against the reference Se4047 genome (Accession number: FM204883) using SMALT v0.7.4 (https://www.sanger.ac.uk/resources/software/smalt). Locations of deletions and short insertions were predicted using pindel (Ye, Schulz, Long, Apweiler and Ning 2009), and then validated by comparing the mapping of reads spanning indels to the reference genome and to a version of the same reference with the predicted indel included. If the inclusion of the indel improved mapping, that indel was retained, and reads realigned around it as per the remapping. SNPs were identified using a combination of samtools (Li, Handsaker, Wysoker, Fennell, Ruan, Homer, Marth, Abecasis and Durbin 2009) mpileup and bcftools as previously described (Harris, Feil, Holden, Quail, Nickerson, Chantratita, Gardete, Tavares, Day, Lindsay et al. 2010). Large deletions and duplications were identified for each isolate based on read coverage along the reference genome using a continuous hidden Markov model with three states: 0x coverage, 1x coverage and ≥2x coverage. Initial and transition frequencies were fitted to the data using a Baum-Welch optimisation, and the most likely sequence of hidden states was calculated using the Viterbi algorithm (Viterbi 1967). IS element insertion locations were identified by mapping read data to a set of IS sequences known to occur in *S. equi* and *S. zooepidemicus*. Where one read of a pair mapped to an IS element, the non-mapping paired read was mapped to the reference Se4047 genome to identify the insertion site. To remove false positive insertion sites that may be caused by chimeric read pairs, a minimum of five reads were required to map to the same insertion location for it to be accepted.

### Phylogenomic Analysis

Maximum likelihood phylogenetic reconstruction of variable sites was performed using RAxML v7.0.3 (Stamatakis 2006) under a general time reversible (GTR) evolutionary model using a gamma correction for among-site rate variation. 100 random bootstrap replicates were run to provide a measure of support for relationships in the maximum likelihood tree. A linear regression of root-to-tip distance versus isolation date was used to assess the fit of a strict molecular clock to the data, and gave a weak correlation for isolates with known isolation dates, excluding vaccine isolates (coefficient of determination, R^2^ = 0.25). To test the null hypothesis that such an R^2^ arose by chance alone, we repeated the root-to-tip analysis 1000 times with the tip dates of the isolates randomly permutated each time (Firth, Kitchen, Shapiro, Suchard, Holmes and Rambaut 2010). In all cases, the data with random permutations gave lower R^2^ values than the real data, so that we could reject our null hypothesis at the 0.001 level and accept the alternative hypothesis that the real data contains significant temporal signal. Bayesian reconstruction in BEAST v1.7 (Drummond and Rambaut 2007b) was used to estimate substitution rates and times for divergences of internal nodes on the tree under a GTR model with a gamma correction for among-site rate variation. All combinations of strict, relaxed lognormal, relaxed exponential and random clock models and constant, exponential, expansion, logistic and skyline population models were evaluated. For each, three independent chains were run for 100 million generations, sampling every 10 generations. On completion each model was checked for convergence, both by checking ESS values were greater than 200 for key parameters, and by checking independent runs had converged on similar results. Models that failed to converge, including logistic population models, were discarded. Models were compared for their fit to the data using Bayes Factors based on the harmonic mean estimator as calculated by the program Tracer v1.4 from the BEAST package. The best-fit model combination was found to be a relaxed exponential clock model and a skyline population model, and so this combination was used for all further analysis. A burn-in of 10 million states was removed from each of the three independent runs of this model before combining the results from those runs with the logcombiner program from the BEAST package. A maximum clade credibility (MCC) tree was created from the resulting combined trees using the treeAnnotator program, also from the BEAST package.

### Accessory Genome Assembly

A pan genome for *S. equi* was created by identifying novel regions from *de novo* assemblies of each isolate. Assemblies were created with Velvet v1.2.09 (Zerbino and Birney 2008) using the VelvetOptimiser.pl v2.2.5 (http://bioinformatics.net.au/software.shtml) script to optimise the kmer length, expected coverage and coverage cut-off parameters based on the N50 statistic. A core genome was defined by removing the mobile prophages, φSeq1-4, and ICE elements, ICE*Se1* and ICE*Se2*, from the Se4047 reference genome. This core genome was then mapped to each assembly using NUCmer (Delcher 2002). All regions >200 bp in each assembly that did not match to the core genome were extracted and retained as accessory regions. All accessory regions for each isolate and the accessory regions from the reference *Se*4047 genome were then mapped against each other using NUCmer. Where two regions were identical in length and matched along their entire length, one was retained. Where one region was completely contained within another the longer region was retained. Any region of novel sequence >200 bp was also retained. Using this process for all pairwise comparisons led to production of a non-redundant set of accessory genome contigs. After filtering, these accessory contigs were appended to the core genome to form a pan genome. Finally, each assembly and the reference *Se*4047 genome were mapped to the pan genome using NUCmer to identify the regions of the pan genome present in each isolate.

### Mutation frequency

Cultures of test bacteria were grown over night in Todd Hewitt (TH) broth at 37 °C in a 5% CO_2_ enriched atmosphere. Cultures were diluted to an OD_600nm_ of 0.5. The number of viable bacteria was enumerated by plating 100 μl of a 10^-6^ dilution onto each of five TH agar plates that were grown at 37 °C in a 5% CO_2_ enriched atmosphere for 24 hours. 100 μl of undiluted culture was plated onto each of five TH agar plates containing 0.03 μg ml^-1^ rifampicin and grown at 37 °C in a 5% CO_2_ enriched atmosphere for 24 hours. Colonies were enumerated, averaged and the resistance frequency was calculated by dividing the total number of colonies by the number of rifampicin resistant colonies. The experiment was repeated in triplicate and an average resistance frequency was calculated across the three experiments. Statistical significance was calculated using an ANOVA test on the whole population.

### Allelic replacement

The generation of the *eqbE* deletion mutant, Δ*eqbE*, has been described previously (Heather, Holden, Steward, Parkhill, Song, Challis, Robinson, Davis-Poynter and Waller 2008).

### Experimental infection of ponies

Ponies were transferred to a containment unit three days before challenge. Each pony was challenged with *S. equi* strain 4047 or the *ΔeqbE* deletion mutant via the spraying of a 2 ml culture containing 5 × 10^7^ cfu into each nostril. Bacteria were grown overnight in Todd Hewitt broth and 10% fetal calf serum (THBS) in a 5% carbon dioxide enriched atmosphere at 37 °C, diluted 40-fold in fresh pre-warmed THBS, further cultivated and harvested at an OD_600nm_ = 0.3. This infection dose has been shown to optimize the infection rate, whilst avoiding overwhelming the host immune response (Guss, Flock, Frykberg, Waller, Robinson, Smith and Flock 2009; Hamilton, Robinson, Sutcliffe, Slater, Maskell, Davis-Poynter, Smith, Waller and Harrington 2006). Ponies were monitored for the onset of clinical signs of disease over a period of three weeks post challenge by daily physical examination, rectal temperature, lymph node swelling and nasal discharge scoring. Blood samples were taken for quantification of fibrinogen and neutrophil levels by total white blood count performed on Beckman-Coulter ACTdiff analyzer with a manual differential count to calculate % neutrophils. Post mortem examination was performed on all ponies following the onset of early clinical signs of infection such as pyrexia and a reluctance to eat dry pelleted food, preferring haylage or water and prior to the onset of later signs according to strict welfare guidelines at the Animal Health Trust, or on reaching the study endpoint at 3 weeks post challenge. The severity of disease pathology was quantified according to a scoring system described previously (Guss, Flock, Frykberg, Waller, Robinson, Smith and Flock 2009).

## DATA ACCESS

Short reads for all sequenced isolates have been submitted to the European Nucleotide Archive (ENA) under the accession numbers listed in Supplementary Table 1.

## ACKNOWLEDGEMENTS

The authors would like to thank the Horserace Betting Levy Board for funding the analysis of the *eqbE* mutant (ref: Vet/prj/758). The Horse Trust funded the collection of isolates from UK outbreaks of strangles (G1606). We are grateful to Prof. John Timoney (University of Kentucky, USA), Prof. Tom Buckley (Irish Equine Centre, Ireland), Nicolas De Brauwere (Redwings Horse Sanctuary, UK), Dr. Jennifer Stewart (Animal Health Trust, UK), Prof. Bengt Guss (Swedish University of Agricultural Sciences, Sweden), Prof. Gunther van Loon (Ghent University, Belgium), Maria Cristina Marotti Campi (Al Khaledia Equine Hospital, Saudi Arabia) and Els Acke (Massey University, New Zealand) for contributing isolates to this study. We thank the core sequencing and informatics teams at the Sanger Institute for their assistance and The Wellcome Trust for its support of the Sanger Institute Pathogen Genomics and Biology groups. SRH, JP and MTGH were supported by Wellcome Trust grant 098051.

### AUTHOR CONTRIBUTIONS

S.R.H. analysed and interpreted the data and wrote the paper. C.R., K.F.S., K.S.W and R.P. performed experiments, carried out data analyses and cultured the isolates. J.P. helped interpret the data and write the paper. M.T.G.H. jointly conceived the project with A.S.W., facilitated sequencing of the isolates, carried out data analysis and helped write the paper. A.S.W. conceived and ran the project, collected samples and helped write the paper.

---

**Supplementary Figure 1.** Representation of predicted homologously-recombined regions. The left panel represents the ML phylogeny of *S. equi*, with BAPs cluster and MLST type shown in columns adjacent to the tree, as in Figure 1. The right panel represents regions identified as exhibiting significantly raised SNP density, indicative of import of variation *en masse* via homologous recombination or regions under high selective pressures. Above the panel is a representation of the genome annotation of *S. equi Se*4047.

**Supplementary Figure 2.** Mutation spectra associated with branches on the tree leading to the outliers in the root-to-tip analysis (Supplementary Fig. 1) and all other branches. ^*^ indicates significant difference to ‘others’ at the 0.1 level, while ^**^ indicates significance at the 0.05 level. Colors correspond to colors in Supp. Fig. 1.

**Supplementary Figure 3.** Mean resistance frequency of long-branch isolates and the reference *Se*4047 *in vitro* to rifampicin, Values represent the means of three independent experiments conducted in triplicate. Error bars indicate 95% confidence intervals.

**Supplementary Figure 4.** Tanglegram showing concordance in BEAST (left) and ML (right) tree topologies, but not branch lengths. Branch lengths in the Bayesian phylogeny produced with BEAST represent time, while those in the ML tree represent genetic diversity. Dates are shown beneath the BEAST tree, and BAPs cluster and MLST type in columns adjacent to the tree.

**Supplementary Figure 5.** Variation in substitution rates between acute and persistent isolates. A) Histogram showing the frequency of branches subtending acute and persistent isolates with different estimated mutation rates. B) Scatter plots of branch length vs mean estimated substitution rate for branches subtending acute, persistent and other (unknown) isolates. The unbroken and dashed red lines in each plot indicate the mean and 95% HPD estimates for the entire data from BEAST.

**Supplementary Figure 6.** Coverage of the accessory genome across the species. The left panel shows the ML tree, with BAPs cluster in a column to the right, as in Figure 1. The right-hand panel shows coverage of the accessory genome in each isolate. To the top and bottom of the right panel are representations of the assembled accessory contigs from isolates in the study. The contig color gives an indication of the content of the contig. Pink = bacteriophage, green = integrative and conjugative element (ICE), blue = transposon, red = IS element. Contigs present in the reference genome are labeled in bold above the panel, along with the location of some important virulence genes. For each isolate in the tree, regions are colored gray if they were present in single copy, and black if they were in multiple copy. i.e. duplications. Missing regions are in white.

**Supplementary Figure 7.** Core genome insertions, duplications and IS elements in persistent, acute and other isolates. Three ML phylogenetic trees are shown in the left panel, created from only persistent isolates (top), only acute isolates (middle) and other isolates (bottom). For each, the column to the right of the tree indicates the BAPs cluster of the isolates in the species phylogeny in Figure 1. In all cases, the topologies of the individual trees are consistent with the tree in Figure 1. The right-hand panel shows coverage of the core genome in each isolate. To the top and bottom are a representation of the annotation of the core genome, with the *has* and *cit* loci labeled. Regions of single copy coverage are colored green in persistent isolates, red in acute isolates and gray in others. Regions of duplication are colored black, and deletions are white. IS element insertion locations are shown in blue.

**Supplementary Figure 8.** Accessory genome insertions, duplications and IS elements in persistent, acute and other isolates. Three ML phylogenetic trees are shown in the left panel, created from only persistent isolates (top), only acute isolates (middle) and other isolates (bottom). For each, the column to the right of the tree indicates the BAPs cluster of the isolates in the species phylogeny in Figure 1. In all cases, the topologies of the individual trees are consistent with the tree in Figure 1. The right-hand panel shows coverage of the accessory genome in each isolate. To the top and bottom of the right panel are representations of the assembled accessory contigs from isolates in the study. The contig color gives an indication of the content of the contig. Pink = bacteriophage, green = integrative and conjugative element (ICE), blue = transposon, red = IS element. Contigs present in the reference genome are labeled in bold above the panel, along with the location of some important virulence genes. Regions of single copy coverage are colored green in persistent isolates, red in acute isolates and gray in others. Regions of duplication are colored black, and deletions are white. IS element insertion locations are shown in blue.

**Supplementary Table 1**. *S. equi* isolates the study. Original Excel spreadsheet available from ftp://ftp.sanger.ac.uk/pub/pathogens/S_equi_Supplementary_Tables

**Supplementary Table 2.** List of variable sites identified in isolates included in the study. Original Excel spreadsheet available from ftp://ftp.sanger.ac.uk/pub/pathogens/S_equi_Supplementary_Tables

